# Lifespan-increasing drug nordihydroguaiaretic acid inhibits p300 and activates autophagy

**DOI:** 10.1101/718833

**Authors:** Tugsan Tezil, Manish Chamoli, Che-Ping Ng, Roman P. Simon, Victoria J. Butler, Manfred Jung, Julie Andersen, Aimee W. Kao, Eric Verdin

## Abstract

Aging is characterized by the progressive loss of physiological function in all organisms. Remarkably, the aging process can be modulated by environmental modifications, including diet and small molecules. The natural compound nordihydroguaiaretic acid (NDGA) robustly increases lifespan in flies and mice, but its mechanism of action remains unclear. Here, we report that NDGA is an inhibitor of the epigenetic regulator p300. We find that NDGA inhibits p300 acetyltransferase activity *in vitro* and suppresses acetylation of a key p300 target in histones (i.e., H3K27) in cells. We use the cellular thermal shift assay to uniquely demonstrate NDGA binding to p300 in cells. Finally, in agreement with recent findings indicating that p300 is a potent blocker of autophagy, we show that NDGA treatment induces autophagy. These findings identify p300 as a novel target of NDGA and provide mechanistic insight into its role in longevity.

## INTRODUCTION

Aging is associated with an increase in age-related diseases that include many cancers, neurodegenerative diseases (e.g., Parkinson’s and Alzheimer’s), atherosclerosis and associated cardiovascular disorders (e.g., heart attacks and stroke), macular degeneration, osteoarthritis and sarcopenia^1^. Remarkably, aging or its manifestations as chronic diseases of aging can be slowed by a number of interventions that include diet, exercise, and pharmaceutical drugs^2^. The Interventions Testing Program at the National Institute on Aging has tested 27 drugs for their effects on lifespan in mice. They identified six drugs, including rapamycin, aspirin, acarbose, nordihydroguaiaretic acid (NDGA), protandim, and 17α-estradiol, that were associated with increased median lifespan^3–6^. In particular, NDGA showed a consistent median lifespan extension by 8–10% at three different doses in mice, particularly in males^5^. Old mice (22 months old) treated with NDGA showed improved grip duration and rotarod performance, indicating better muscle function than untreated age-matched animals^6^. NDGA also extends median lifespan in evolutionarily distant organisms, such as fruit flies (by 12%) and mosquitoes (by 50%)^7,8^. Several studies also showed the effects of NDGA on neurodegenerative disorders. In a fruit fly model of Parkinson’s disease, NDGA delayed the loss of climbing ability associated with onset of neurodegeneration^9^. NDGA also decreased motor dysfunction in a mouse model of amyotrophic lateral sclerosis and extended lifespan by 10% in this context ^10^. Furthermore, NDGA restored synapse structure and extended lifespan by 19% in a mouse model of Huntington’s disease^11^ and decreased amyloid-beta deposition in the brains of mouse models of Alzheimer’s disease^12,13^.

*In vitro* studies show that NDGA scavenges hydroxyl radical, peroxynitrite, superoxide anion and singlet oxygen, suggesting that NDGA might function as an antioxidant agent^14,15^. *In vivo*, NDGA exerts cytoprotective effects by modulating the nuclear factor erythroid 2-related factor 2 (Nrf2)/antioxidant response element (ARE) antioxidant pathway^16^. This variety of effects makes NDGA a compelling candidate for further study in the context of aging and geroprotective processes.

The hallmarks of aging include epigenetic alterations and deregulation of nutrient sensing^1^. Levels of histone acetylation change during aging and are related to increased acetyl-CoA levels in aging animals^17^, and acetyl-CoA levels decrease significantly upon calorie restriction even before cellular ATP, NADH or amino acid levels are affected^18,19^. As acetyl-CoA is the only acetyl-group donor for protein acetylation reactions, its cellular level highly influences the activity of acetyltransferases^20^. Particularly, histone and non-histone acetyltransferase p300 (E1A-associated protein p300) functions as a molecular sensor of changes in cellular and possibly other acyl-CoA levels as well^21^. Any decrease in cellular acetyl-CoA significantly limits the activity of p300 and decreases subsequent acetylation reactions involving key cellular proteins^22,23^. Notably, p300 potently blocks autophagy by acetylating autophagy-related proteins including Atg5, Atg7, Atg12 and LC3^24^.

Although the health benefits of NDGA have been demonstrated in different organisms and conditions, its underlying mechanism of action remains unclear. Early reports show that NDGA inhibits arachidonic acid 5-lipoxygenase^25^, which catalyzes the first step in leukotriene A4 biosynthesis, and lysophosphatidylcholine acyltransferase-2 (LPCAT2)^26^, an enzyme in platelet-activating factor (PAF) biosynthesis that converts the precursor lyso-PAF (1-O-alkyl-sn-glycero-3-phosphocholine) into active PAF (1-O-alkyl-2-acetyl-sn-glycero-3-phosphocholine) by acetylation. Interestingly, salicylate (an active metabolite of aspirin) and diflunisal (a salicylate derivative) both also inhibit LPCAT2^27^.

We recently reported that both salicylate and diflunisal are inhibitors of the histone acetyltransferase p300^28^. Here we tested the possibility that NDGA also inhibits p300 and exerts its effect on aging by inhibition of this important epigenetic regulator. We report that NDGA inhibits p300. We further present evidence of NDGA directly binding to p300 in cells and show that NDGA treatment activates the autophagy pathway and promotes autophagosome formation in human cells, as well as in *C. elegans*.

## RESULTS

### NDGA inhibits p300 *in vitro*

To test the hypothesis that NDGA is a p300 inhibitor, we used two different *in vitro* enzymatic assays that measure the acetyltransferase activities of p300 and PCAF (p300/CBP-associated factor). In the first assay, we used purified human recombinant p300 BHC protein^29^ (a construct containing the bromodomain, catalytic acetyltransferase domain, and C/H3 domain, amino acids 965–1810) that represents the acetyltransferase activity of full length protein^30^, histone H3 (amino acids 1–21) peptide as a substrate, and acetyl-CoA as an acetyl-group donor, to measure the activity of p300 on histone H3 in the presence of NDGA. In this fluorimetric assay, NDGA inhibited p300 BHC with an IC_50_ of 10.3 μM, compared to 75 μM for the known p300 inhibitor anacardic acid^31^ (Fig. 1A). When we evaluated NDGA for the inhibition of human recombinant PCAF HAT (histone acetyltransferase domain), we observed no inhibition even at high concentrations of NDGA (Fig. 1B), ruling out a nonspecific mechanism of action. In the second assay, we used purified human recombinant p300 HAT protein consisting of the core histone acetyltransferase domain only (amino acids 1284–1673), histone H3 peptide (amino acids 1–21) and acetyl-CoA mixture in reaction buffer containing increasing concentrations of NDGA. Consistent with Fig. 1A, B, NDGA inhibited p300 HAT activity (Fig. 1C) but failed to inhibit purified human PCAF HAT activity (Fig. 1D). To further understand the mechanism of p300 inhibition, we determined inhibition kinetics of p300 HAT activity by NDGA for both substrates acetyl-CoA and H3 peptide. Double-reciprocal plots were constructed to find the nature of inhibition. Inhibition kinetics studies clearly showed that NDGA is a noncompetitive inhibitor for both acetyl-CoA (Fig. 1E) and histone H3 binding sites (Fig. 1F). These results indicate that NDGA is a selective inhibitor of p300 acetyltransferase activity *in vitro*.

**Fig. 1.**
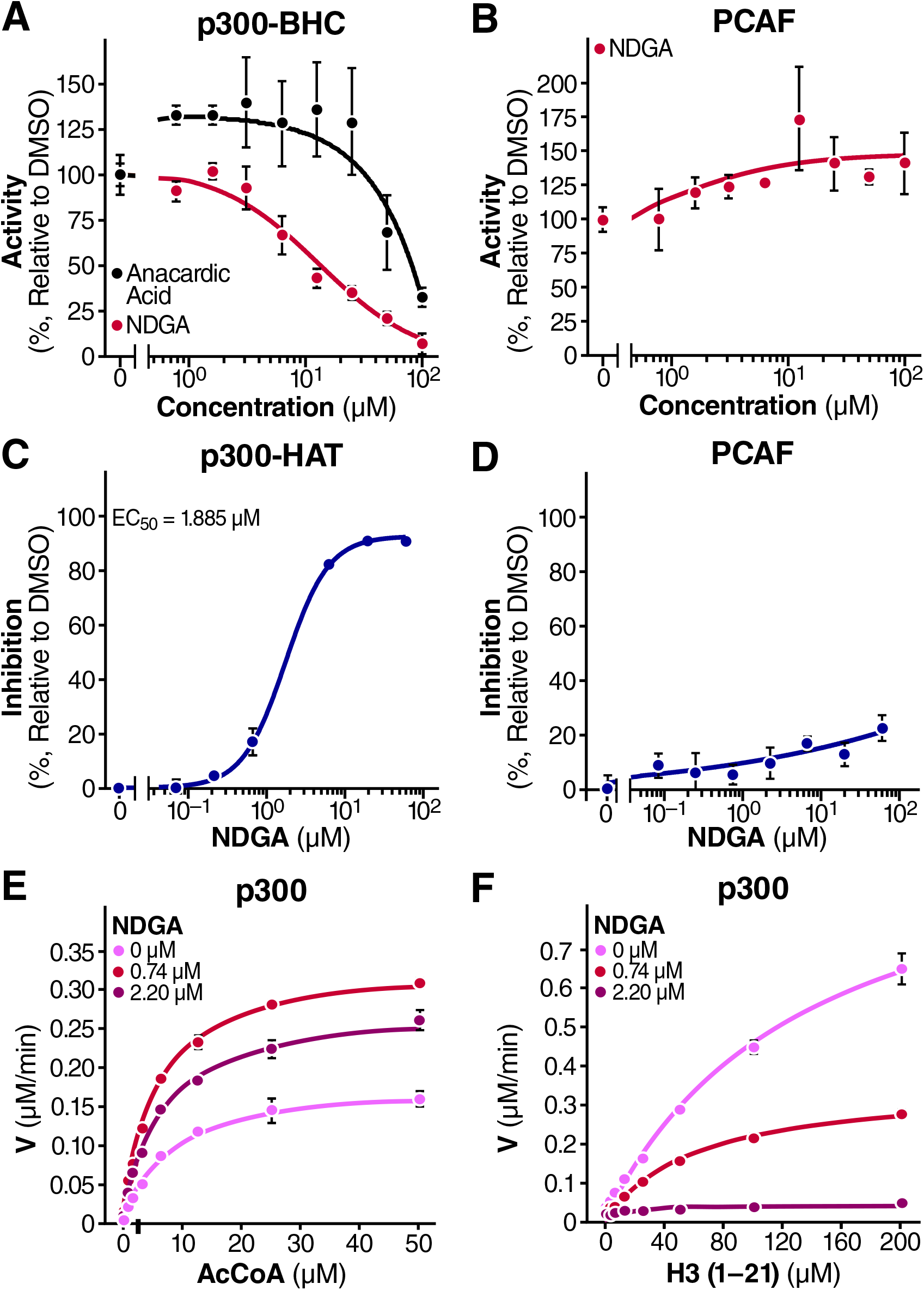
NDGA inhibits histone acetyltransferase activity of p300 *in vitro*. **A.** The acetyltransferase activity of p300 on histone H3 peptide (aa 1–21) was measured in the presence of NDGA or anacardic acid. Results are expressed as percentage activity in the absence of NDGA. A paired t-test was performed for statistical significance, *p* = 0.0013. **B.** Effect of NDGA on PCAF HAT activity expressed as percentage activity in the absence of NDGA. **C, D**. Inhibitory effects of NDGA on p300 (panel C) and PCAF (panel D) using another enzymatic assay were measured and are expressed as percentage inhibition relative to control DMSO. **E, F**. Effects of varying concentrations of acetyl-CoA (panel E) and substrate peptide histone H3 in the presence of varying concentrations of NDGA were measured (panel F). Data represent the results from three independent experiments (mean ± S.D.), and curves were generated using nonlinear regression fit.

### NDGA inhibits p300-mediated histone acetylation in human, mouse and fruit fly cells

As an epigenetic regulator, p300 orchestrates gene expression via acetylation of specific lysine residues in specific cellular histone proteins in cells^32^. To determine whether NDGA influences histone acetyl marks in cells, we monitored the acetylation status of different lysine residues in HEK293T cells treated with NDGA. Specifically, we extracted the histones and conducted western blot analysis using antibodies selective for unique acetylated histone residues and determined IC_50_ values for each residue (Fig. 2A, Suppl. Fig. 1). Previous genetic studies reported that deletion of p300 specifically reduces acetylation on H3K27 but not H3K9^33^. Consistent with our hypothesis that NDGA selectively inhibits p300 in cells, we found that histone H3K27 acetylation was the most suppressed in cells treated with NDGA with an IC_50_ of 8.8 μM (Fig. 2A-C). The acetylation of other histone residues was also suppressed but in most cases with IC_50_ >25 μM, except for H3K14 (Fig. 2A). As expected, residues that are not p300 targets, such as H3K9, were unchanged by NDGA (Fig. 2A-C).

**Fig. 2.**
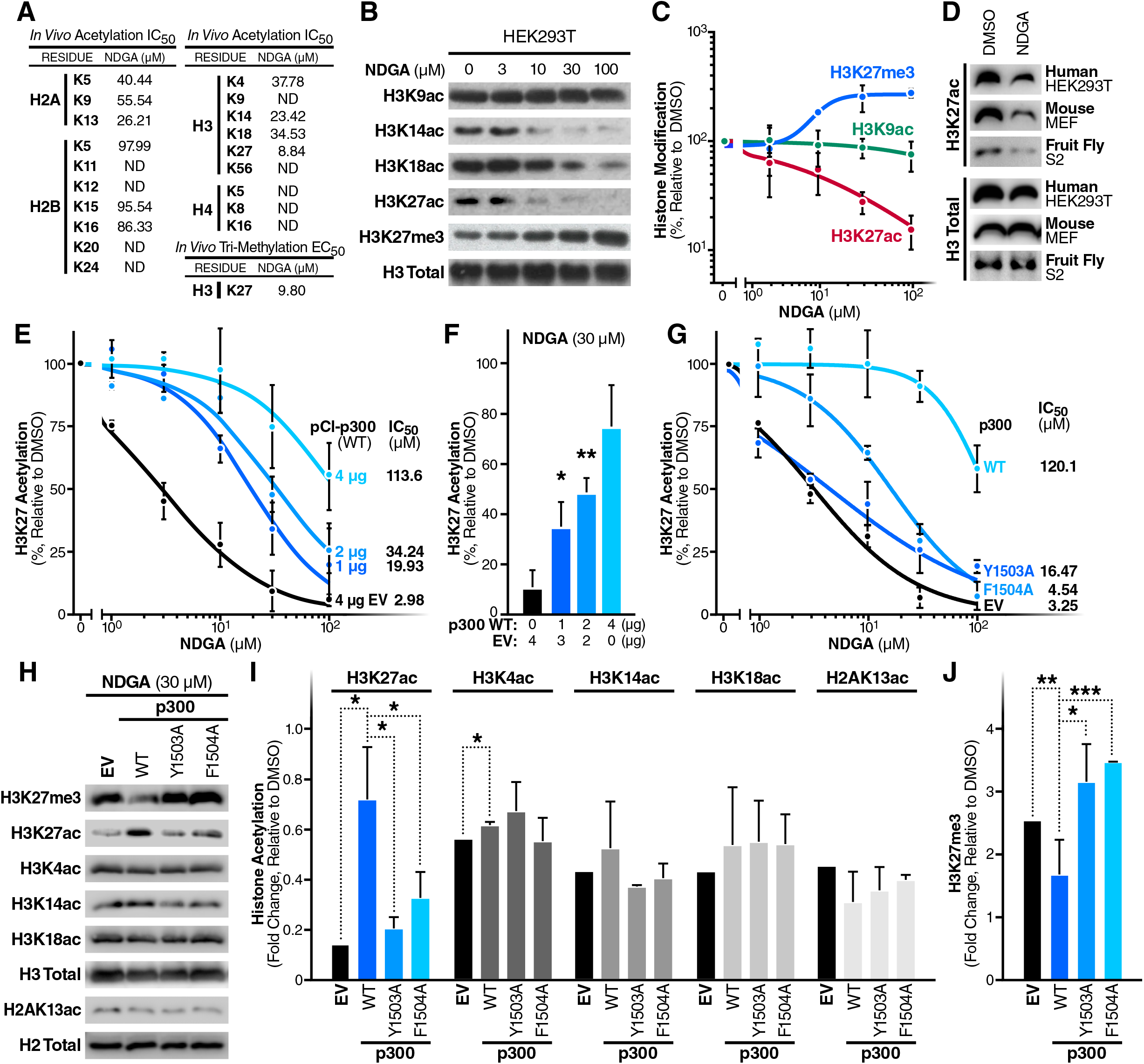
NDGA inhibits p300-driven histone acetylation in HEK293T cells. **A.** Median inhibitory concentration (IC_50_) values for NDGA on different acetylated histone residues and median effective concentration (EC_50_) value on H3K27 tri-methylation were determined by immunoblotting of histones extracted from cells after 24-hour treatments with varying concentrations of NDGA (ND, not determined). **B.** Differences in acetylation and tri-methylation of p300 target (H3K14, H3K18, H3K27) and non-target (H3K9) histones were monitored by immunoblotting after indicated concentrations of NDGA treatment for 24 hours and histone extraction. **C.** Dose-dependent changes in acetylation and tri-methylation on H3K27 after 24-hour NDGA treatment were analyzed by immunoblotting and are represented as percentage change in histone modification relative to DMSO treatment. **D.** Decreases in H3K27 acetylation upon 30 μM of NDGA treatment for 24 hours in human (HEK293T), mouse (MEF) and fruit fly (S2 Schneider) cells were monitored by immunoblotting after histone extraction. **E.** Effects of wildtype p300 overexpression on NDGA-dependent H3K27 hypoacetylation were determined by densitometric analysis of immunoblotting data. Cells were transfected with different amounts of human wildtype EP300 encoding expression plasmid pCI-p300(WT) and/or empty vector (EV) and treated with varying NDGA concentrations for 24 hours. The data represent percentage change in H3K27 acetylation relative to DMSO treatment and the change in IC_50_ values. **F.** The rescue of p300 overexpression on H3K27 hypoacetylation after 30 μM NDGA treatment was performed as in panel E. **G.** Effects of overexpression of wildtype and catalytically inactive p300 mutants (Y1503A and F1504A) on NDGA-induced H3K27 hypoacetylation were determined by immunoblotting and densitometry. Cells were transfected by the same amount of expression plasmids or empty vector (EV) and treated with varying NDGA concentrations. Histones were extracted and analyzed. **H.** The effects of wildtype and mutant (Y1503A or F1504A) p300 overexpression on NDGA-induced histone hypoacetylation and hypermethylation were determined by immunoblotting after 30 μM NDGA treatment for 24 hours. **I.** Effects of wildtype and mutant p300 overexpression on NDGA-induced histone hypoacetylation were analyzed by immunoblotting and densitometry. **J.** Decreased H3K27 tri-methylation upon wildtype p300 overexpression was detected by immunoblotting and densitometry. Total histone antibodies were used as loading controls in all experiments. Data represent the results from two independent experiments (mean ± S.D.) and curves were generated using nonlinear regression fit.

To test the effect of NDGA on H3K27 acetylation in different organisms, we also treated mouse (MEF) and fruit fly (S2 Schneider) cell lines with NDGA (30 μM) for 24 hours, and analyzed extracted histones by immunoblotting. As with HEK293T, H3K27 acetylation was also found to be decreased also in these cell lines upon NDGA treatment (Fig. 2D, Suppl. Fig. 2), demonstrating the suppressive effect of NDGA on H3K27 acetylation in evolutionarily distant organisms.

### NDGA-induced suppression of H3K27ac is associated with increased tri-methylation of the same residue in HEK293T cells

Decreased histone H3 K27 tri-methylation (H3K27me3) is a characteristic hallmark of aging in both invertebrates and vertebrates^1^. Worms and fruit flies exhibit an age-associated decrease in H3K27me3 marks^34^, but in humans, a strong decrease in this modification has been observed in primary fibroblasts from patients with Hutchinson-Gilford progeria syndrome, a premature aging disease^35^. Conversely, the naked mole rat, which is the longest-living rodent with a lifespan of more than 28 years^36^, bears higher levels of H3K27 tri-methylation compared to mice^37^. Since acetylation and methylation are mutually exclusive on a single lysine, decreased acetylation can result in an increase in methylation on the same residue^38^. Therefore, we treated HEK293T cells with increasing concentrations of NDGA, extracted histones, and determined the methylation status of H3K27 by immunoblotting. As predicted and in agreement with the decrease in H3K27 acetylation, cells treated with NDGA showed a significant increase in H3K27 tri-methylation with a half maximal effective concentration (EC_50_) of 9.80 μM (Fig. 2A-C).

### Overexpression of p300 reverses NDGA-induced loss of H3K27 acetylation

To further assess p300 as a relevant target of NDGA, we overexpressed different amounts of full-length p300 in HEK293T cells and treated them with NDGA. In support of a role for p300 in sustaining H3K27 acetylation, we found that overexpression of p300 suppressed the effect of NDGA in a dose-dependent manner and increased H3K27 acetylation (Fig. 2E, Suppl. Fig. 3). The IC_50_ of NDGA in rescue experiments was strongly correlated with the amount of p300-expressing plasmid transfected (Fig. 2F and Suppl. Table). Furthermore, addition of catalytically inactive p300 mutants (p300Y1503A and F1504A)^39^ failed to revert NDGA-mediated inhibition (Fig. 2G, Suppl. Fig. 3). In addition to H3K27 acetylation, we also evaluated the acetylation of other histone residues with an IC_50_ lower than 50 μM (H3K4, H3K14, H3K18, H2AK13). While we observed a significant rescue effect on H3K27 and a slight rescue in H3K4 acetylation with wildtype p300 overexpression, changes in other acetylated residues were insignificant (Fig. 2H-I, Suppl. Fig. 4). In agreement with our previous result, we observed a significant decrease in H3K27 tri-methylation when wildtype p300 was overexpressed; whereas overexpression of mutant p300 was unable to change the levels of H3K27me3 (Fig. 2J, Suppl. Fig. 4). These findings support the hypothesis that NDGA-mediated H3K27 hypoacetylation and elevated H3K27 tri-methylation are specifically due to inhibition of p300 acetyltransferase activity.

### p300 is a direct target of NDGA in HEK293T cells

To provide additional evidence that NDGA directly binds to p300 in cells, we used a cellular thermal shift assay (CETSA) that uniquely allows monitoring of drug engagement inside cells^40^. Based on the principle of ligand-induced protein stabilization, compound-bound proteins precipitate at higher temperatures than unbound proteins and can be detected at higher levels in the soluble fraction. To identify direct targets of NDGA, we tested different acetyltransferases. We exogenously expressed p300 HAT (p300/CBP family), GCN5 (general control of amino acid synthesis protein 5-like 2) and PCAF (GNAT family), and TIP60 (60-kDa Tat-interactive protein, MYST family) in HEK293T cells and treated the cells with NDGA. Intact cells were heated at different temperatures for 3 minutes to denature and precipitate proteins. These cells were then subjected to three freeze-thaw cycles to lyse them, isolation of a soluble fraction after centrifugation (20,000 x g) and analysis by immunoblotting. We observed an increase in the thermostability of p300 HAT domain by 10.8°C upon NDGA treatment (Fig. 3A, Suppl. Fig. 5). To confirm this thermostabilization in a dose-dependent manner, we treated cells with different concentrations of NDGA and heated them at a single temperature (55°C) where we previously detected 35% of the total amount of p300 in the soluble fraction. Maximal stabilization (100%) occurred at 30 μM NDGA (Fig. 3B, Suppl. Fig. 5). Importantly, CETSA analysis of GCN5, PCAF and TIP6 (Fig. 3C-E, Suppl. Fig. 5) showed no significant change in thermostability upon NDGA treatment. The difference in melting temperatures (T_m_) of proteins tested are shown in Fig. 3F. These results further support the model that NDGA specifically binds to p300 HAT at a concentration consistent with its inhibitory activity on histone acetylation (Fig. 2).

**Fig. 3.**
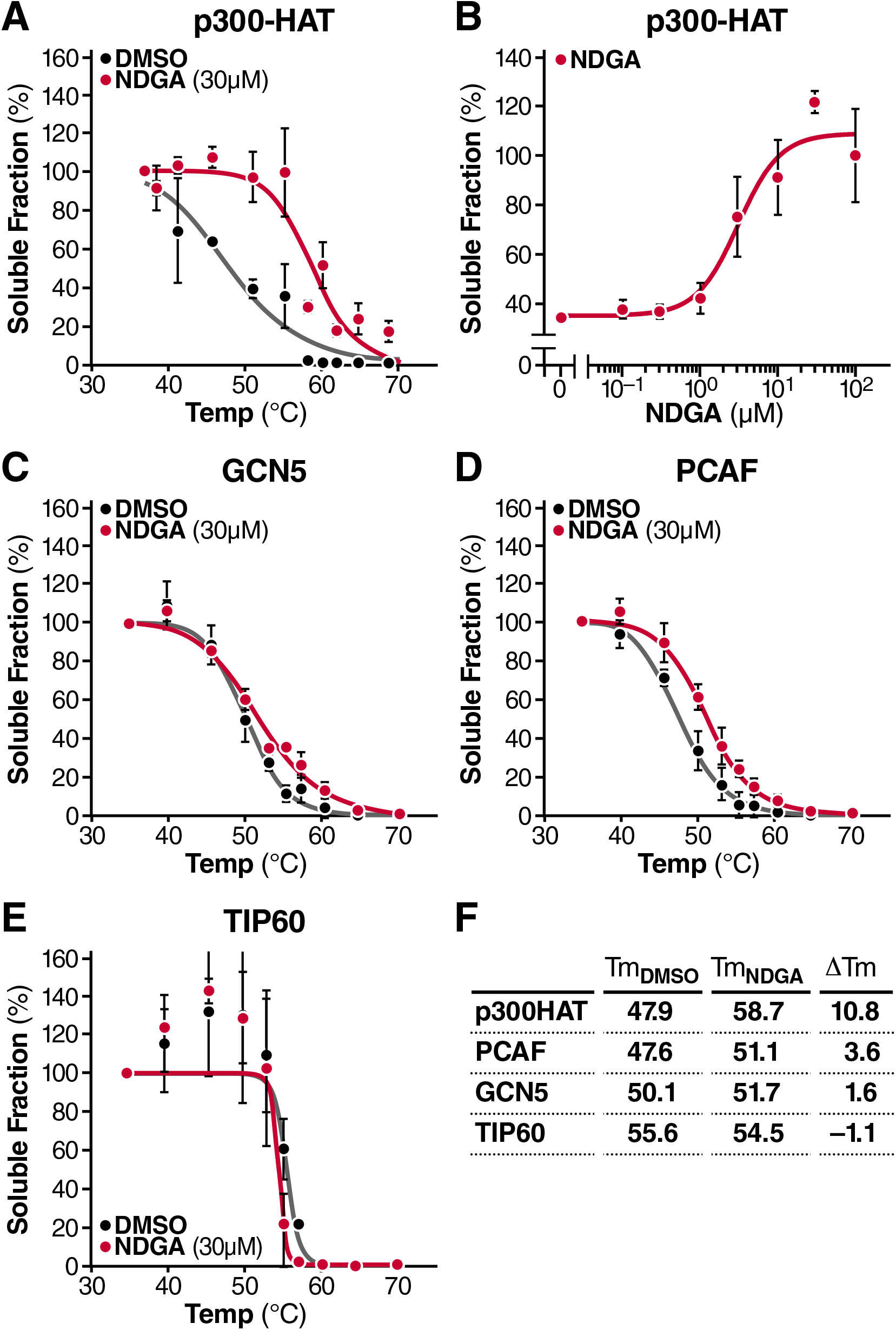
NDGA is a target of NDGA in HEK293T cells. **A.** Melting curves of the p300 HAT domain were generated from a CETSA experiment performed in intact cells expressing HA tagged p300-HAT construct following 3-hour DMSO or NDGA treatment. **B.** Isothermal dose-response fingerprint-CETSA (ITDRF_CETSA_) graph was generated from cells expressing the p300-HAT construct. Intact cells were treated with different concentrations of NDGA for 3 hours and subjected to heating at 55°C for 3 minutes to monitor thermostabilization. **C-E.** Melting curves of different acetyltransferases were generated from intact cells expressing HA-GCN5 (panel C), HA-PCAF (panel D) or HA-TIP60 (panel E) after 3-hour DMSO or 30 μM NDGA treatment and CETSA. **F.** Change in melting temperatures (T_m_) of tested proteins upon NDGA treatment. Data represent the results from two independent experiments (mean ± S.D.), and curves were generated using nonlinear regression fit.

### NDGA induces autophagy in HEK293T and HeLa cells

While we demonstrated a potential molecular mechanism for the role of NDGA in aging cells above, the cellular mechanism by which NDGA elicits these effects remains unexplored. p300, the molecular target of NDGA, is a regulator of cellular autophagy, an evolutionarily conserved process for efficient removal of damaged and potentially harmful cellular contents, including long-lived proteins and cellular organelles^41^. To accomplish this cellular cleansing effort, the coordinated actions of various autophagy-related (Atg) proteins are required. Atg proteins provide the main molecular machinery essential for initiating autophagosome formation via an ubiquitin-like conjugation system^42^. Interestingly, p300 modulates the acetylation status of several Atg proteins. Silencing of p300 expression reduces the acetylation of Atg5, Atg7, LC3 and Atg12, and increases their stability and cellular level, resulting in autophagy pathway activation. Overexpression of p300, on the other hand, causes a significant increase in acetylation, decreases the stability of mentioned key proteins and consequently blocks autophagy by lowering their cellular level^24^. These previous results predict that inhibiting p300 by NDGA could act through activating autophagy. To test this hypothesis, we measured the level of autophagy after NDGA treatment and in response to serum-free media as a positive control. Adding NDGA to HEK293T cells induced several molecular markers of autophagy, including AMPK phospho-activation and upregulation of Beclin-1 (Fig. 4A, Suppl. Fig. 6).

**Fig. 4.**
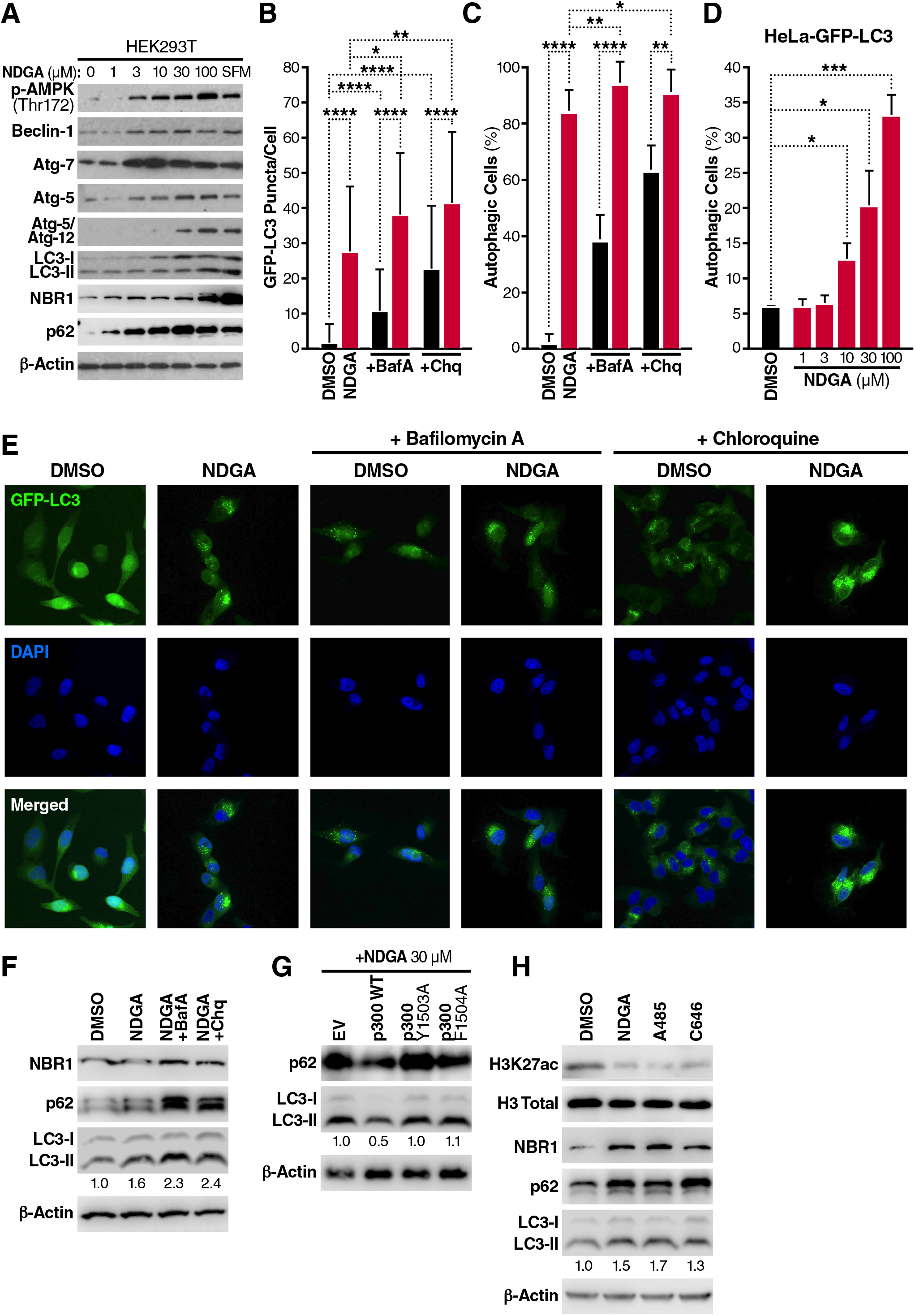
NDGA induces autophagy in HEK293T and HeLa cells. **A.** Activation of the autophagy signaling pathway was monitored by immunoblotting of total proteins isolated from HEK293T cells after a 24-hour treatment with DMSO or varying concentrations of NDGA. Serum-free medium (SFM) was used as a positive control of autophagy. **B.** Autophagosome formation in HeLa-GFP-LC3 cells was monitored by counting GFP-LC3 puncta after a 24-hour treatment with DMSO or NDGA, with or without Bafilomycin A1 (BafA, 100 nM) or Chloroquine (Chq, 100 μM) treatment 3 hours before sample collection. **C.** The percentage of autophagy was measured in HeLa-GFP-LC3 cells treated with DMSO or NDGA, with or without Bafilomycin A1 or Chloroquine treatment for 3 hours prior to sample collection. The threshold value to determine autophagy-positive cells was based on previous measurements and DMSO-treated cells; **D.** Autophagy induction in HeLa-GFP-LC3 cells was analyzed by FACS after a 24-hour treatment with varying NDGA concentrations. **E.** GFP-LC3 puncta (green) in HeLa-GFP-LC3 cells treated with DMSO or NDGA, with or without Bafilomycin A1 or Chloroquine were captured by confocal microscopy. Cells were fixed after 24-hour treatment, stained with DAPI (blue) and analyzed. **F.** Effects of Bafilomycin A and Chloroquine treatment for 3 hours on autophagy marker proteins were quantitated by immunoblotting following total protein extraction. **G.** Effects of wildtype and mutant (Y1503A and F1504A) p300 overexpression on NDGA-induced autophagy were monitored by immunoblotting. **H.** Effects of p300 inhibitors on H3K27 acetylation and autophagy induction were analyzed by immunoblotting. HEK293T cells treated with NDGA (100 μM), A485 (100 nM) or C646 (100 μM) for 24 hours, and total protein or histones were extracted. Numbers in immunoblotting images (F-H) indicate the fold change determined by densitometry analysis. β-Actin and total histone H3 were used as loading controls. Data represent the results from two independent experiments (mean ± S.D.), and statistical significance was determined by student t-test (\**p*<0.05; \*\**p*<0.01, \*\*\**p*<0.001, \*\*\*\**p*<0.0001).

Atg proteins together regulate the formation of a double membrane structure (autophagosome) that engulfs the cellular cargo targeted for degradation^43^. Briefly, Atg7 induces Atg12-Atg5 conjugation and this conjugate specifies localization of LC3 on the lipid membrane. Our results showed a dose-dependent increase in Atg7 levels, and an increase in Atg5/Atg12 dimerization (Fig. 4A, Suppl. Fig. 6), consistent with induction of autophagy. Additionally, we observed upregulation of LC3-II, which is associated with autophagosome formation as well as upregulation of the autophagy cargo receptors NBR1 (Neighbor of BRCA1 gene 1) and p62 (SQSTM1, Sequestosome-1) upon treatment with NDGA (Fig. 4A, Suppl. Fig. 6). To confirm the induction of autophagy in cells, we visualized the process by imaging HeLa cells stably expressing GFP-LC3. Following a 24-hour treatment with NDGA (30 μM) or DMSO, we analyzed LC3 lipidation. LC3-positive puncta were significantly increased with NDGA treatment. Moreover, treatment with Bafilomycin A1, an V-ATPase inhibitor that prevents the fusion of autophagosomes and lysosomes^44^, or with the lysosomal pH-neutralizing compound Chloroquine^45^ resulted in an additional increase in GFP-LC3 puncta in treated cells (Fig. 4B and Fig. 4E). These results indicate the accumulation of autophagosomes demonstrating that HeLA-GFP-LC3 cells have a functional autophagy pathway. An increase in autophagosome number upon NDGA treatment caused a rise in the number of cells undergoing autophagy (Fig. 4C and Fig. 4E). We also observed that the pro-autophagic effect of NDGA was dose-dependent (Fig. 4D, Suppl. Fig. 7). NDGA-treated HEK293T cells showed increased levels of endogenous NBR1, p62 and LC3-II upon NDGA treatment; Bafilomycin A or Chloroquine treatment further enhanced this effect (Fig. 4F, Suppl. Fig. 6). Since the overexpression of p300 rescues H3K27 hypoacetylation (Fig. 2F), we next evaluated p62 and LC3-II levels in wildtype versus mutant p300 overexpressing HEK293T cells upon NDGA treatment. Fig. 4G shows the rescue effect of excess p300 expression by downregulation of these autophagy markers (Suppl. Fig. 6). When we compared NDGA with other p300 inhibitors C646 and A485 for autophagy induction, we discovered that these inhibitors also increased the levels of autophagy markers NBR1, p62 and LC3-II in HEK293T cells (Fig. 4H, Suppl. Fig. 6). These data indicate that NDGA effectively increases autophagic flux in HEK293T and GFP-LC3 HeLa cells, and that p300 activity is a key element in NDGA-induced autophagy.

### NDGA increases median lifespan, decreases histone acetylation, and induces autophagy in worms

To determine the prolongevity property of NDGA *in vivo*, we treated wildtype *C. elegans* (N2 Bristol) with 100 μM NDGA. The median lifespan of worms exposed to NDGA was 21.4% greater than that of worms exposed to the vehicle alone (DMSO) (Fig. 5A). To rule out potential variations in food availability caused by NDGA, we examined the effect of NDGA on feeding bacteria (OP50) and found that OP50 growth was unaffected by NDGA (Fig. 5B). These findings indicate that NDGA has a beneficial effect on *C. elegans* lifespan that is independent of the feeding bacteria growth.

**Fig. 5.**
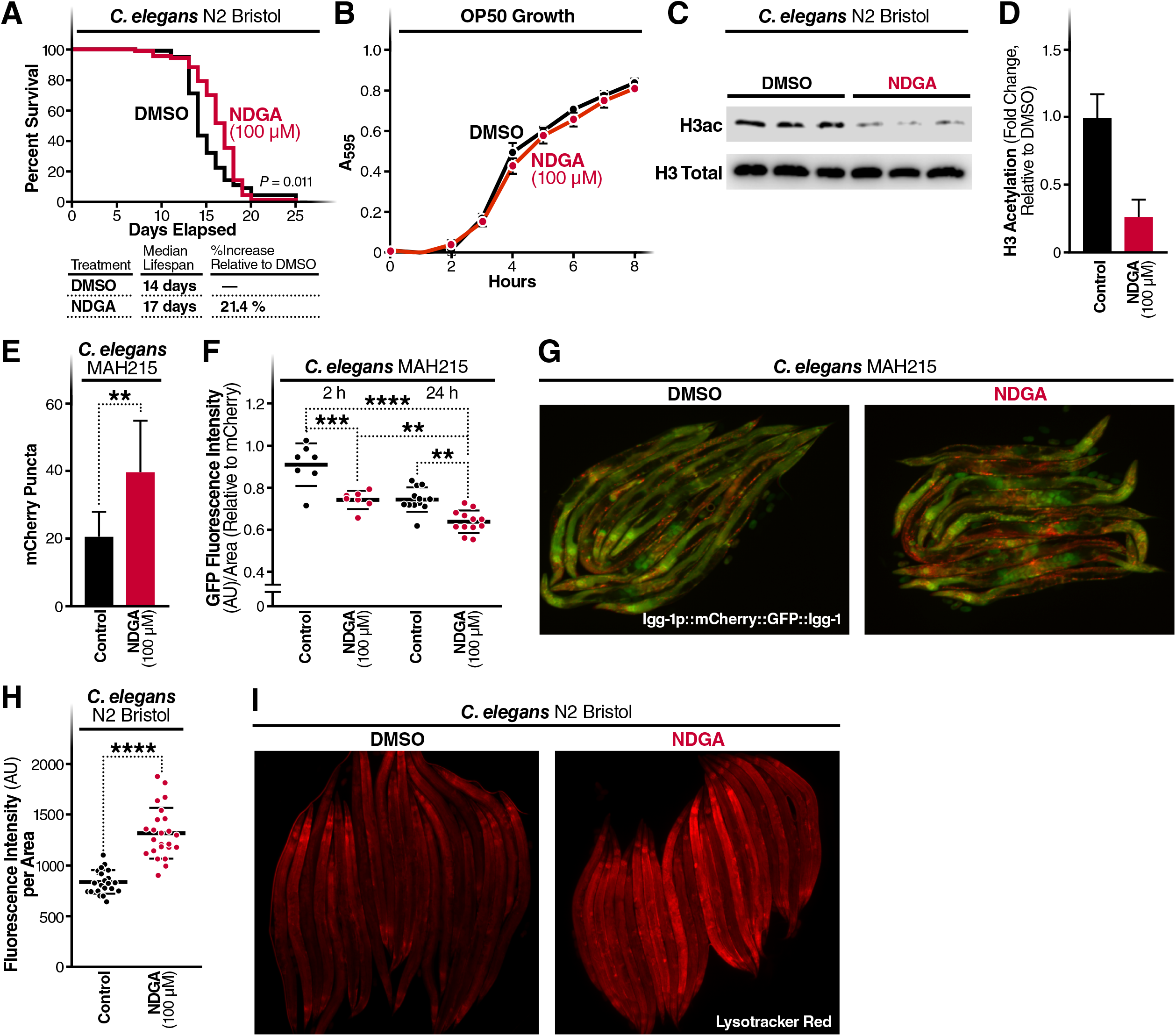
NDGA increases median lifespan and induces autophagy in worms. **A.** The effect of NDGA on worm lifespan was measured by exposing N2 worms to DMSO or NDGA (100 μM) on NGM plates. Average values from three independent experiments were plotted as a lifespan curve. Lower panel indicates the median lifespan of each group as days and percentage; **B.** The effect of NDGA on feeding bacteria (*E. coli* OP50) growth was measured spectrophotometrically (A_595_) every hour for 8 hours in LB medium with 100 μM of NDGA or DMSO. **C and D.** Total histone H3 acetylation in worms was measured by immunoblotting after 24-hour NDGA (100 μM) or DMSO exposure and total protein extraction. **E.** Changes in mCherry puncta number in MAH215 worms were monitored by fluorescence microscopy after a 24-hour NDGA or DMSO treatment. **F.** Effects of NDGA treatment on GFP by time were measured by fluorescence microscopy after NDGA or DMSO (Control) treatment for 2 and 24 hours and represented relative to mCherry intensity. **G.** Monitoring of tandem reporter *lgg-1p::mCherry::GFP::lgg-1* expressing worms was performed by fluorescence microscopy. **H and I.** Monitoring Lysotracker Red staining in N2 worms after 24-hour treatment with NDGA (100 μM) or DMSO was performed by fluorescence microscopy, and fluorescence intensity was measured for individual animals in each group. Data represent the results from three independent experiments (mean ± S.D.), and statistical significance was determined by student t-test (\**p*<0.05; \*\**p*<0.01, \*\*\**p*<0.001).

CBP-1, the nematode orthologue of p300/CBP, is responsible for histone acetylation in *C. elegans*, and animals with catalytically inactive cbp-1 have decreased global histone acetylation^46^. With this knowledge, we evaluated the possible suppressive effect of NDGA on histone H3 acetylation. We treated worms with 100 μM NDGA for 24 hours, analyzed histone H3 acetylation by immunoblotting and observed a significant decrease in histone H3 acetylation in treated animals (Fig. 5C-5D, Suppl. Fig. 8).

To analyze autophagy induction in worms in depth, we used described reporter strain (MAH215) expressing pH sensitive GFP in a tandem *lgg-1p::mCherry::GFP::lgg-1* in which mCherry puncta indicates autophagosomes and autolysosomes^47^. Worms exposed to NDGA had an increase in mCherry-only puncta, indicating an elevated autophagic activity as assessed by increased fusion of autophagosomes to lysosomes (Fig. 5E and 5G). MAH215 worms with this tandem reporter allow us to analyze the autophagosome maturation process by monitoring the GFP and mCherry fluorescence intensity over time. Increased autophagic flux and lysosomal activity following NDGA treatment is also evident from observed decreases in pH-sensitive GFP fluorescence over time resulting from the quenching of GFP within the acidic lysosome. We examined the NDGA-treated worms (2–24 hours) by measuring the ratio of GFP relative to mCherry fluorescence. We observed that GFP signal was significantly less in animals treated for 2 hours than in the vehicle-treated animals (DMSO), and those levels continued to decrease at 24 hours (Fig. 5F). In agreement with these results, worms treated with NDGA also show increased lysotracker staining, suggesting an increased number of lysosomes/late endosomes most likely as a result of increased autophagic activity (Fig. 5H, 5I). These data provide evidence that NDGA treatment causes an increase in autophagic flux in *C. elegans*.

## DISCUSSION

Here we present evidence for a novel potential mechanism for NDGA actions as an lifespan increasing drug. We showed that NDGA effectively inhibits p300 histone acetyltransferase activity *in vitro*, and it directly binds the p300 HAT domain and alters several epigenetic marks associated with aging in human, mouse and fruit fly cells. Notably, while we identify p300 as a target of NDGA, we cannot exclude the possibility that it blocks the activity of other acetyltransferases. For example, CREBBP (CBP), which shares high levels of homology with the p300 catalytic domain, is likely to be also targeted by NDGA. But importantly, NDGA failed to enhance the thermostability of GNAT family members PCAF and GCN5 and the MYST family member TIP60 in CETSA experiments.

p300/CBP is conserved in different organisms from nematodes to mammals and plays an essential role in cell and organism viability. Worms with inactive CBP-1, knockout of dCBP (the fruit fly homolog of p300/CBP), and homozygous p300 or CBP knockout mice demonstrate embryonic lethal phenotypes^46,48,49^. These observations indicate that the use of potent pharmacologic inhibitors of p300 in clinic might cause unintended toxicity in younger animals. Therefore, p300 inhibition may be more applicable during late adulthood or for the treatment of age-related diseases that rely on elevated p300 activity such as cancer and neurodegenerative diseases^50,51^. Several chemical inhibitors with varying potency and selectivity for p300 have already been tested for their potential anti-cancer properties, including anacardic acid^52^, C646^53^ and A485^54^.

Previous studies on NDGA carried out with mice fed diets containing different doses of NDGA have determined that blood levels of NDGA in supplemented animals (165 nM) are lower than those we found to be required to inhibit p300 in cell-free systems and to activate autophagy in cells^5^. However, it should be noted that tissue and plasma concentrations of a compound are not linearly correlated factors. In fact, several studies previously reported this phenomenon^55,56^. The concentration of NDGA in serum, uptake in different tissues and its intracellular concentration may vary. Currently, it is unknown how NDGA is metabolized, which proteins are involved in its cellular uptake in different tissues, and how NDGA is distributed in the body. These characteristics of NDGA should be further investigated in different cells and tissues.

Previous studies on mice indicate a sexual dimorphism in response to NDGA treatment: males showed an increase in median lifespan, but females did not^5^. Importantly, this study also showed that the median lifespan of control female mice is approximately 10% higher than males, and that NDGA-treated males demonstrate a level of benefit similar to the control lifespan curves of females. Therefore, NDGA may provide beneficial effects in both females and males, but possible adverse effects in females could prevent lifespan extension. Moreover, several compounds extend lifespan differently favoring either sex. For instance, acarbose and 17-α estradiol show either smaller or no lifespan extension in female mice, compared to males^5^. Aspirin and rapamycin inhibit inflammatory response, which could have a sex-dependent effect in late-life inflammation^57^. Sex difference in autophagy induction has also been documented and may influence the efficacy of a compound^58^. The dimorphic nature in response to NDGA or other lifespan-extending compounds still needs to be understood with further studies.

Activation of autophagy has emerged as an important intervention against aging and the chronic diseases of aging, and its inhibition reverses the beneficial effects of anti-aging interventions in all species investigated to date^59^. Importantly, p300 decreases the stability of Atg5, Atg7, Atg12 and LC3 by acetylation and functions as an anti-autophagic regulator under high nutrient conditions^24^. Our findings are in accord with several reports showing autophagy induction upon inhibition of p300^60^. Our results highlight the role of p300 and pharmacological inhibitors, such as NDGA, in key aging-related molecular and cellular processes. These observations support the hypothesis that the suppressor function of p300 on autophagy can also be regulated pharmacologically.

Moreover, other interventions that increase lifespan, such as spermidine, aspirin, calorie restriction and rapamycin, also provide their beneficial effects in part by inducing autophagy, potentially through p300 inhibition61-64. While spermidine and aspirin directly inhibit p300, calorie restriction diminishes p300 activity indirectly by lowering cellular acetyl-CoA levels23,65. Rapamycin indirectly inhibits p300 by suppressing mTORC1, which phosphorylates p300 and prevents its intra-molecular inhibition, resulting in autophagy suppression66. Together with our results, these findings underscore the significant role of p300 in aging and could generate renewed interest in evaluating the clinical utility of p300 inhibitors for both aging and age-related diseases.

## METHODS

### Cell culture

HEK293T (ATCC), HeLa-LC3-GFP and MEF (ATCC) cells were maintained in appropriate volume of Dulbecco’s Modified Eagle Medium supplemented with 10% serum (Serum Plus - II, Sigma), 100 units/mL penicillin and 100 units/mL streptomycin (Corning, VA). Schneider’s *Drosophila* Line 2 (S2 Schneider) was kindly provided by Henri Jasper, Buck Institute for Research for Aging. S2 cells were grown at 25°C in Schneider’s Insect Medium (Himedia) that was prepared according to the manufacturer’s instructions.

### Chemicals and buffers

NDGA (Sigma), C646 (Sigma), A485 (Tocris) and Bafilomycin A1 (Adipogen) were dissolved in dimethyl sulfoxide (DMSO) (Sigma-Aldrich), Chloroquine (Biovision) was dissolved in water. Phosphate-buffered saline (PBS) was purchased from Corning. Tris-buffered saline with Tween (TBST) buffer (150 mM NaCl, 0.01% (v/v) Tween-20, 50 mM Tris-HCl buffer, pH 7.6) was used for immunoblotting. Blocking buffer and primary antibody incubation solution was 5% (w/v) BSA (Sigma), pH 7.0, in TBST. Secondary antibody incubation buffer was 5% (w/v) blocking reagent (Sigma) in TBST.

### Plasmids and transfection

HEK293T cells were transfected using TransIT-X2 (Mirus) reagent, following the manufacturer’s instructions, and further treatments were carried out 48 hours post transfection. Full-length wildtype p300, p300(Y1503A) and p300(F1504A)^28^ were used for rescue experiments. Primers used for cloning are listed in Supplementary Information.

### *In vitro* acetyltransferase activity assays

The activities of histone acetyltransferases, p300 and PCAF, were measured using a fluorimetric assay kit, following the manufacturer’s instructions (SensoLyte p300 Assay kit, SensoLyte PCAF Assay kit, Anaspec) using appropriate dilutions of NDGA and anacardic acid. To measure the inhibition of p300 and PCAF, appropriate dilutions of NDGA and anacardic acid were prepared and added to the assay mixture. The manual *in vitro* acetyltransferase assay kit was performed in reaction buffer (final concentration; 40 mM HEPES pH 7.5, 40 mM NaCl, 0.025 *%* Triton X-100, 50 μM histone H3 peptide (amino acids 1–21), 40 nM p300 (Enzo Life Science, BML-SE451) or 30 nM PCAF (Cayman Chemical, 10009115)) in black 96-well microtiter plates (PerkinElmer, 6005320). Appropriate dilutions of NDGA were prepared and added to the assay mixture. Enzyme concentrations were evaluated to give a linear response between enzyme concentration and initial velocity for each acetyltransferase tested. Negative controls received assay buffer without peptide. Mixtures were allowed to equilibrate for 5 minutes at room temperature, enzymatic reactions were initiated by addition of acetyl-CoA (50 μM final concentration) and incubated for 20 minutes at 37 °C. Upon completion, reactions were stopped with 1:1 volume of cold isopropanol and CPM (7-diethylamino-3-(4’-maleimidylphenyl)-4-methylcoumarin, 12.5 μM final concentration). After incubation for 15 minutes at room temperature, plates were read in a fluorescence microplate reader (POLARStar OPTIMA, BMG Labtech) at Ex = 390 nm / Em = 460 nm with a gain setting of 1391. For competition analysis, concentrations of acetyl-CoA or histone H3 peptide were varied over 0.78–50 μM and 3.1–200 μM, respectively. Standard curves were generated by preparing a twofold dilution series of freshly dissolved acetyl-CoA in assay buffer covering a concentration range of 0–50 μM. Samples were then mixed with isopropanol, reacted with CPM, and measured as described above.

### SDS-PAGE and immunoblotting

HEK293T cells were treated with NDGA for 24 h and lysed in lysis buffer (20 mM Tris-HCl pH 7.5, 150 mM NaCl, 1 mM Na2EDTA, and 1 mM EGTA, 1% TritonX100) supplemented with complete EDTA-free protease inhibitor cocktail (ROCHE) and PhosStop phosphatase inhibitor cocktail (ROCHE). Polyacrylamide gels (10–15%) were used for the separation of total proteins. 30 μg of total proteins were transferred to 0.2 μm nitrocellulose membranes using the BioRad blotting system. Histones were extracted from HEK293T cells by EpiQuik Total Histone Extraction Kit (Epigentek), following manufacturer’s instructions. 1–5 μg of histone isolates were separated by Polyacrylamide gels (15%) and transferred to 0.5-μm nitrocellulose membranes using the BioRad blotting system. All membranes were blocked with blocking buffer (5% BSA in TBST) and probed with primary and then secondary antibody. Antibodies used are shown in Supplementary Information. Lumi-Light Western Blotting Substrate (ROCHE) or SuperSignal West Femto Maximum Sensitivity Substrate (Thermo Fisher) were used for detection of the target proteins. Chemiluminescence intensities were detected using the ChemiDoc imaging system (BioRad) and quantified using Image J software (NIH). β-actin levels were used to normalize the intensities of bands. For CETSA curves, the band intensities were related to the intensities of the lowest temperature for the control samples or drug-exposed sample. For the ITDRF_CETSA_ experiments, the band intensities were related to control samples. All experiments were performed at least twice. All blots derive from the same experiment and were processed in parallel.

### Cellular thermal shift assay (CETSA)

For CETSA, HEK293T cells were freshly seeded the day before the experiment. The day of experiment, equal numbers of cells were counted by Moxi Mini automated cell counter (Orflo), and 0.6 × 10^6 cells per temperature were seeded in T-25 cell culture flasks (VWR) in an appropriate volume of culture medium. Cells were exposed to 100 μM NDGA or equal volume DMSO for 3 hours in an incubator with 5% CO_2_ and 37°C. After the incubation, cells were harvested, washed with PBS, and resuspended in PBS supplemented with EDTA-free complete protease inhibitor cocktail (ROCHE). Intact cells were divided into 100-μl aliquots and heated individually at different temperatures for 3 minutes in a PCR machine (Thermal cycler, BIO-RAD), followed by cooling for 2 minutes at room temperature. Cell suspensions were freeze-thawed three times using liquid nitrogen, and the soluble fraction was separated from the cell debris by centrifugation at 20000 x g for 20 minutes at 4°C. Supernatants were transferred to new micro centrifuge tubes and analyzed by SDS-PAGE, followed by immunoblotting analysis. Immunoblotting results were subjected to densitometry analysis (ImageJ, NIH), and melting temperatures of p300-HAT, PCAF, GCN5 and TIP60 were determined by Prism 7 software (GraphPad) using nonlinear regression fit. The viability of the cells was assessed in triplicate by trypan blue exclusion. All CETSA experiments were performed at least as triplicates.

Isothermal dose response fingerprint-cellular thermal shift assay (ITDRF_CETSA_) HEK293T cells were treated with different concentrations of NDGA and prepared as described above. Intact cells were aliquoted (0.6 × 10^6 cells per concentration) into PCR tubes and heated at 55°C, where 35% of p300 HAT protein remained in the soluble fraction) for 3 minutes. Samples were cooled down for an additional 2 minutes at room temperature and freeze-thawed three times using liquid nitrogen. The soluble fraction was separated by centrifugation at 20000 x g for 20 minutes at 4°C and subjected to immunoblotting. ITDRF_CETSA_ experiments were performed at least as triplicates.

### Flow cytometry

HeLa GFP-LC3 cells were analyzed by FACS as described^67^. Briefly, cells were collected by trypsin and washed with cold PBS. The pellets were resuspended with 0.05% Saponin (diluted in PBS), vortexed briefly and added six volumes of PBS to further dilute the detergent. Cells were pelleted, resuspended with PBS and monitored by FACS (BD Bioscience). Results were analyzed by FACSDiva 8.0.2 software.

### Confocal microscopy

GFP-LC3 HeLa cells were seeded onto glass slide-chamber (Watson Bio Lab) and treated with NDGA (30 μM) for 24 hours and prepared for confocal microscopy. Cells were fixed with 1.6% paraformaldehyde (Alfa Aesar) for 15 minutes at RT and washed with PBS. Then, to stain nuclei, GFP-LC3 HeLa cells were incubated with DAPI (Genetex) for 5 minutes at RT. After three PBS washes, slides were prepared with mounting solution Tris-MWL 4-88 (Citifluor) for imaging. Samples were viewed on a temperature controlled Zeiss 780 LSM confocal fluorescent microscope (488 nm for GFP, 405 nm for DAPI, ×63/1.4 oil-immersion objective) in the Buck Institute for Research on Aging Morphology Core. 16-bit images (2048×2048, C:2, Z:1, T:1) were analyzed using Zen 2.3 2011 software (Zeiss) and Image J software (NIH). GFP-LC3 puncta-positive cells were quantified using the Image J object counter plug-in. Cells that contained 15 or more puncta per cell were considered LC3-positive, and this threshold value (15 puncta/cell) was used to determine the percentage of autophagic cells as described^68^.

### *Caenorhabditis elegans* maintenance and lifespan assay

*C. elegans* hermaphrodites were maintained on nematode growth medium (NGM) agar plates seeded with *E. coli* bacterial strain OP50 at 20°C as described^69^. Strains used in this study were wildtype Bristol N2 and MAH215 (sqIs11 [lgg-1p::mCherry::GFP::lgg-1 + rol-6] rollers^47^. Wildtype N2 Bristol worms were synchronized by a 2-hour egg lay from gravid adult hermaphrodites, and the eggs were transferred to NGM plates containing 100 μM NDGA with a bacterial OP50 lawn at 20°C. Worms were scored as dead when they no longer responded to gentle prodding with a platinum wire. The lifespan of an individual was defined as the time elapsed from when it was placed onto the fresh NGM plate (t=0) to when it was scored as dead. Nematodes that crawled off plates during the assay were excluded from calculations.

### OP50 growth assay

*E. coli* OP50 culture was grown in LB overnight at 37°C shaker (200 rpm). The day after, the culture was diluted 1:1000 in LB containing either 100 μM NDGA or the same volume of DMSO. The absorbance at 595 nm was measured by SpectraMax i3X every hour. Experiments were performed in eight technical replicates in a total of three independent experiments and plotted as time-dependent growth curve.

### Mounting worms and imaging

NDGA (100 μM) was prepared in sterile water and added to the top of NGM plates seeded with a bacterial OP50 lawn. On their first day of adulthood, synchronized population of worms were transferred to either DMSO control or NDGA-treated plates for experimental analysis. Microscopic slides containing pads of 2% agarose (Sigma) were prepared 30 minutes before mounting worms. Around 2–5 μl of 2 mM levamisole (Sigma) was pipetted out on the center of the agarose pads. Worms were transferred to the levamisole drop, and a cover slip was placed on top before imaging.

### Lysotracker red staining

Synchronized day-1 adult N2 worms were transferred to DMSO control or NDGA-treated plates for 24 hours at 20°C. On day 2, worms were transferred to OP50-seeded plates treated with NDGA and containing 1 μM LysoTracker Red DND-99 (Molecular Probes) for 24 hours at 20°C. After 24 hours, worms were picked and washed in a fresh drop of 1X M9 medium and mounted on 2% agarose pad slides for microscopic visualization. Worm images were taken in a Zeiss Imager Z1 fluorescence microscope using rhodamine filters. Image analysis was performed by measuring mean LysoTracker fluorescence of each worm using Image J (NIH) software.

### Tandem-tagged LGG-1 reporter imaging

MAH215 worms were synchronized and transferred to DMSO control or NDGA-treated plates on day 1 of adulthood as above. After 2 or 24 hours, worms were imaged using GFP and rhodamine filters on a Zeiss fluorescence microscope. For each worm, we quantified the number of mCherry-only puncta (i.e., autolysosmes (AL) (the total number of mCherry-positive punctae – the number of GFP-intensity) as described^47^.

### Worm protein extraction

Synchronized day-1 adult N2 worms were transferred to DMSO control or NDGA treated plates for 24 hours at 20°C. 100 worms from each treatment condition were collected in 1X M9 solution. Worms were washed twice and suspended in RIPA lysis buffer. Worm pellets were sonicated for 10 cycles at maximum intensity in Bioruptor sonicator (Diagenode). Lysate was centrifuged at 1000 g for 1 minute, and clear protein lysate was used for immunoblotting.

### Statistical analysis

Dose-response analysis of histone acetylation in cells was analyzed by nonlinear regression fit to Y = 100/(1 + 10 (Log IC50 – X)*H), where H = Hill slope (variable). IC_50_ values represent the concentration of NDGA that inhibits 50% of the occurrence of acetylation of the target residue. The experiments in which wildtype and mutant p300 plasmids used were subjected to correlation analysis by a Pearson correlation. CETSA data were expressed as means ± SD. Band intensities from independent experiments (n ≥ 2) were plotted using nonlinear regression fit (variable slope). ITDRF_CETSA_ data were fitted using a sigmoidal (variable slope) curve fit. Multiple confocal microscopy images (n ≥100 cells per condition) were analyzed, and data were expressed as means ± SD. P values were determined by unpaired, two-tailed Student’s t tests. (\**p* < 0.05; \*\**p* < 0.01; \*\*\**p* < 0.001; **** *p* < 0.0001). P values in lifespan curves were calculated using the log-rank (Mantel-Cox) test. In the tandem-tagged LGG-1 reporter imaging, the average puncta were calculated, and data were analyzed using one-way analysis of variance ANOVA. All calculations were performed using Prism 7 (GraphPad) software.

## DATA AVAILIBILITY

The datasets generated during and/or analyzed during the current study are available from the corresponding author on reasonable request.

## ACKNOWLEDGMENTS

We thank Davalyn Powell for her helpful discussions during the preparation of the manuscript and editorial assistance; John Carroll for preparation of the figures. This study was supported by Glenn Center Postdoctoral Fellowship (T.T.) and support from the Buck Institute for Research on Aging. M.C. is supported by Larry L. Hillblom Foundation. R.P.S. and M.J. have been supported by the Deutsche Forschungsgemeinschaft (DFG, RTG1976).

## AUTHOR CONTRIBUTIONS

T.T. designed and performed experiments, analyzed data and wrote the manuscript. C.N. performed *in vitro* studies in Fig. 1A and B and was responsible from the provision of study materials; R.S. conducted assays in Fig. 1C-F with the supervision of M.J.; M.C. conducted assays in Fig. 5E-I with the supervision of J.A.; T.T. conducted assays in Fig. 5A and B together with V.J.B. under the supervision of A.W.K.; E.V. supervised the study and edited the manuscript. All authors discussed the results and contributed to the final manuscript.

