## Supplementary Information for "Lifespan-increasing drug nordihydroguaiaretic acid inhibits p300 and activates autophagy"

**Supplementary Information Inventory**

**Supplementary Figure Legends**

**Supplementary Figures 1 – 8**

**Supplementary Table**

**Supplementary Notes**

**Supplementary Figure Legends**

**Supplementary Fig. 1.** Uncropped blots corresponding to Figure 2A-C. HEK293T cells were treated with different concentrations of NDGA or DMSO for 24 hours and isolated histones were subjected to immunoblotting for different acetylated histone residues as indicated. Antibodies against total histones were used as loading controls. Data represent at least two independent experiments.

**Supplementary Fig. 2.** Uncropped blots corresponding to Figure 2D.

**Supplementary Fig. 3. A.** Uncropped blots corresponding to Figure 2E-F. HEK293T cells were transfected with different amounts of full-length wildtype pCI-p300 plasmid (0–4 µg of plasmid DNA/well in 12-well plate) and treated with DMSO or varying concentrations of NDGA for 24 hours. Isolated histones were subjected to immunoblotting to detect acetylated histone H3 K27. **B.** Uncropped blots corresponding to Figure 2G. HEK293T cells were transfected with 4 µg of wildtype or catalytically inactive p300 encoding plasmid DNA (pCI-p300Y1503A and pCI-p300F1504A) in a 12-well plate and treated with DMSO or indicated concentrations of NDGA for 24 hours. Isolated histones were monitored for histone H3 K27 acetylation. Total histone H3 was used as a loading control. Data represent at least two independent experiments.

**Supplementary Fig. 4.** Uncropped blots corresponding to Figure 2H-J.

**Supplementary Fig. 5. A.** Uncropped blot of p300-HAT corresponding to Figure 3A. **B.** Uncropped blots of p300-HAT in soluble and insoluble fractions after isothermal dose-response fingerprint-CETSA ($ITDRF_{CETSA}$) at 55°C corresponding to Figure 2B. **C-E.** Uncropped blots of GCN5, PCAF, and TIP60 corresponding to Figure 3 C-E.

**Supplementary Fig. 6. A.** Uncropped blots corresponding to Figure 4A. **B.** Uncropped blots corresponding to Figure 4F. **C.** Uncropped blots corresponding to Figure 4. **D.**

**Supplementary Fig. 7.** Flow cytometry images corresponding to Figure 4D. Cells were permeabilized by saponin (0.05%) and swollen cells were analyzed for insoluble GFP-LC3 signal.

**Supplementary Fig. 8.** Uncropped blots corresponding to Figure 5 C-D.

**Supplementary Figure 1**

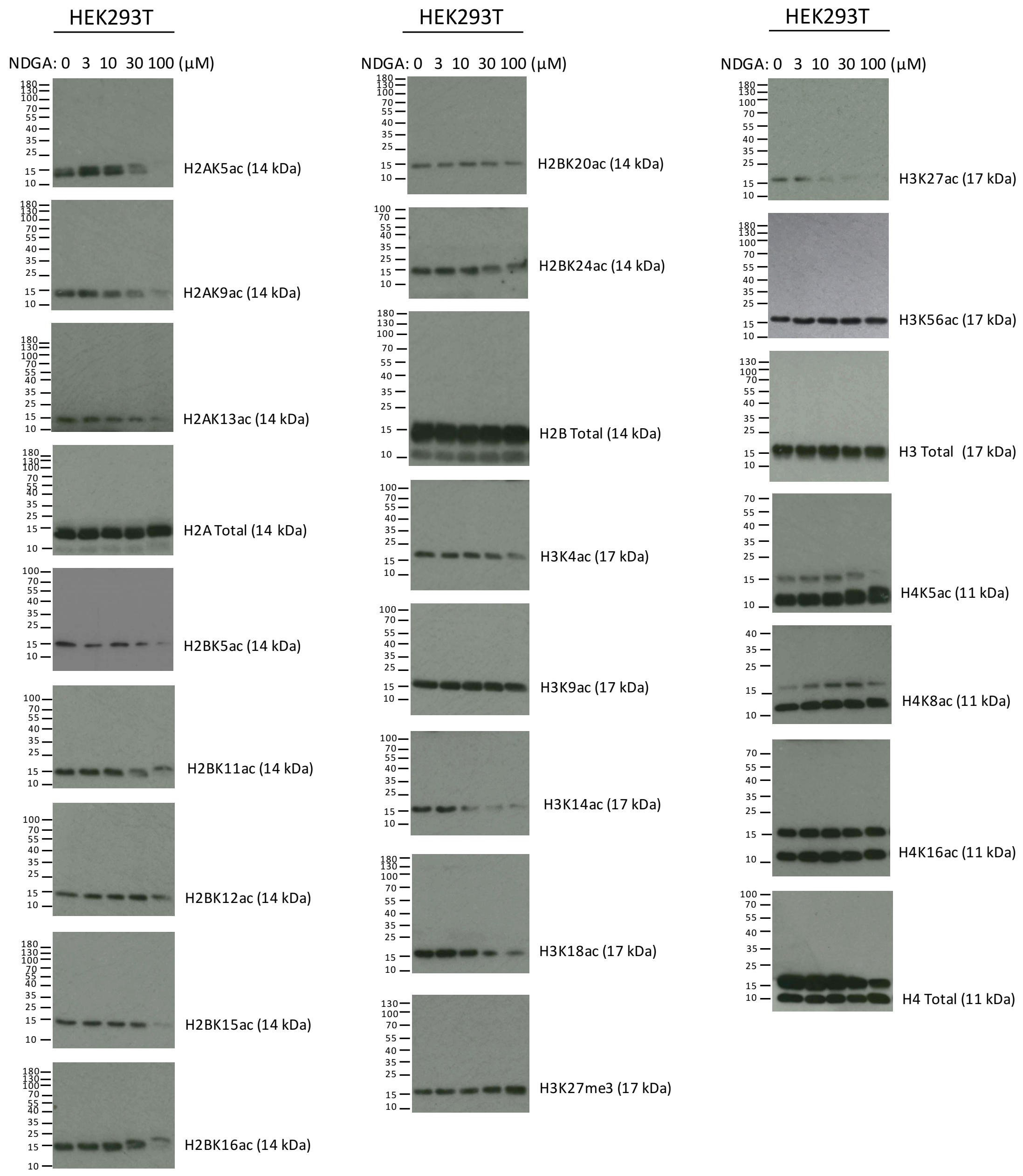

#### Supplementary Figure 2

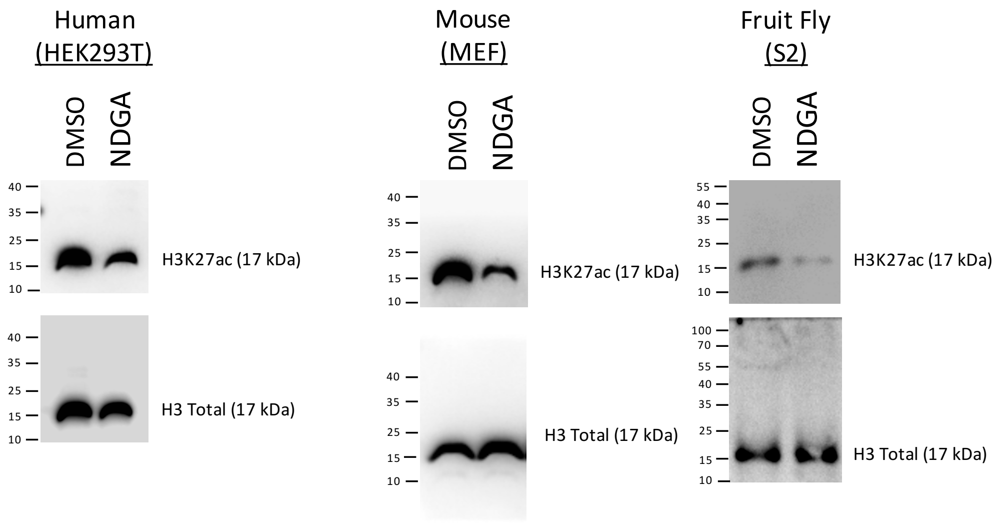

**Supplementary Figure 3**

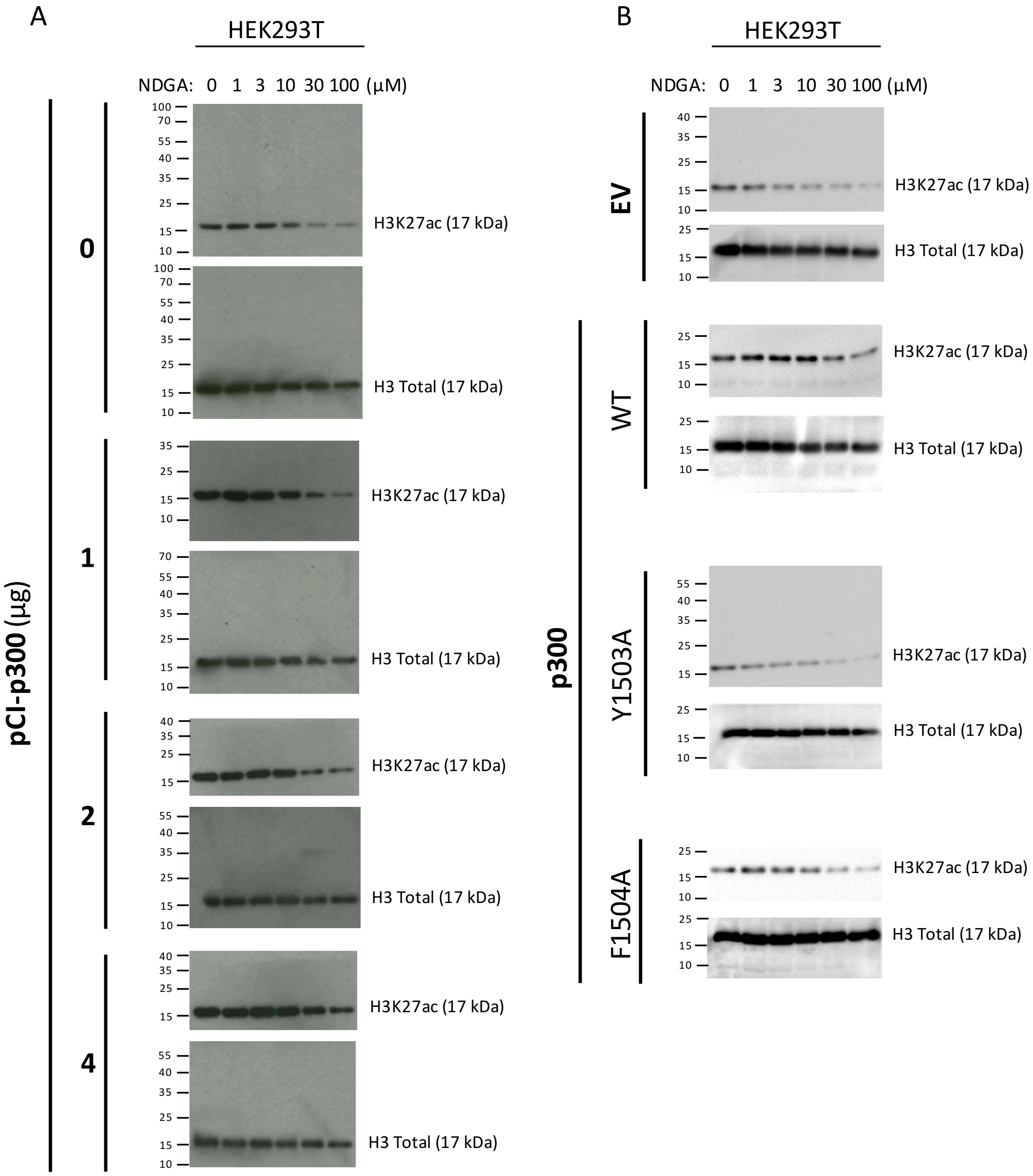

### Supplementary Figure 4

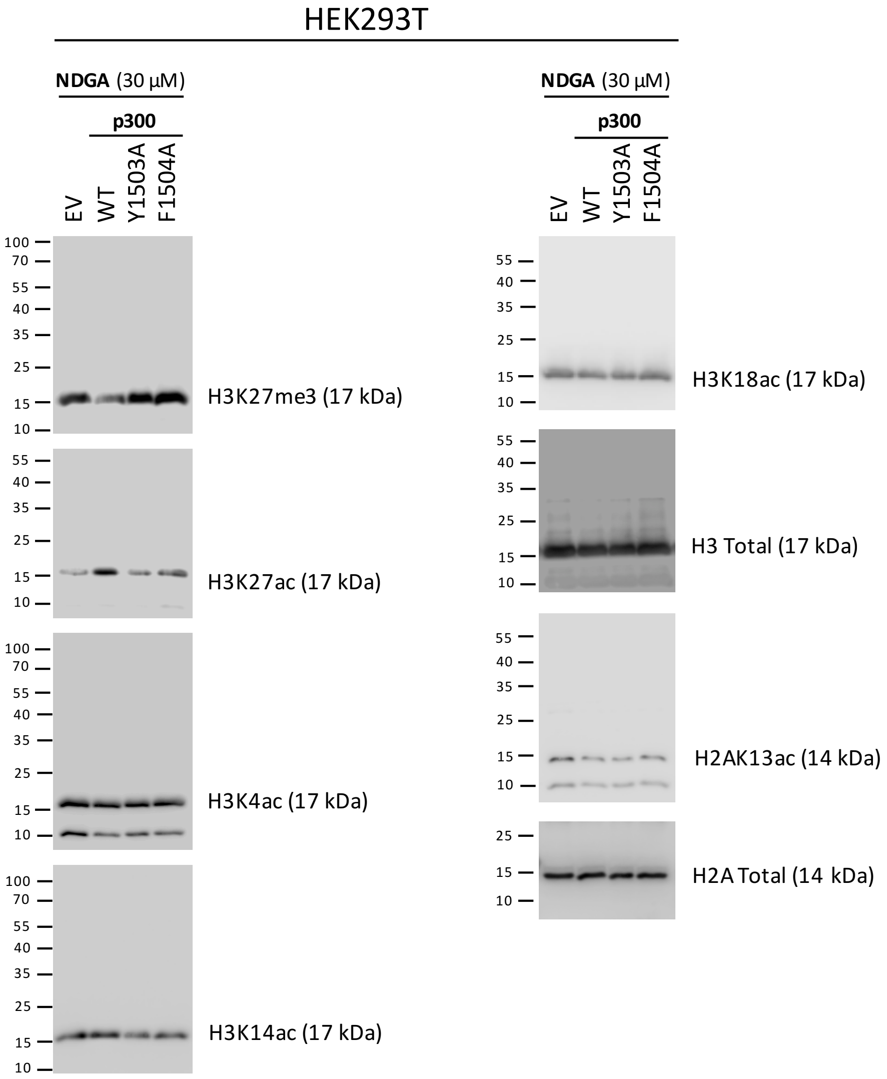

#### Supplementary Figure 5

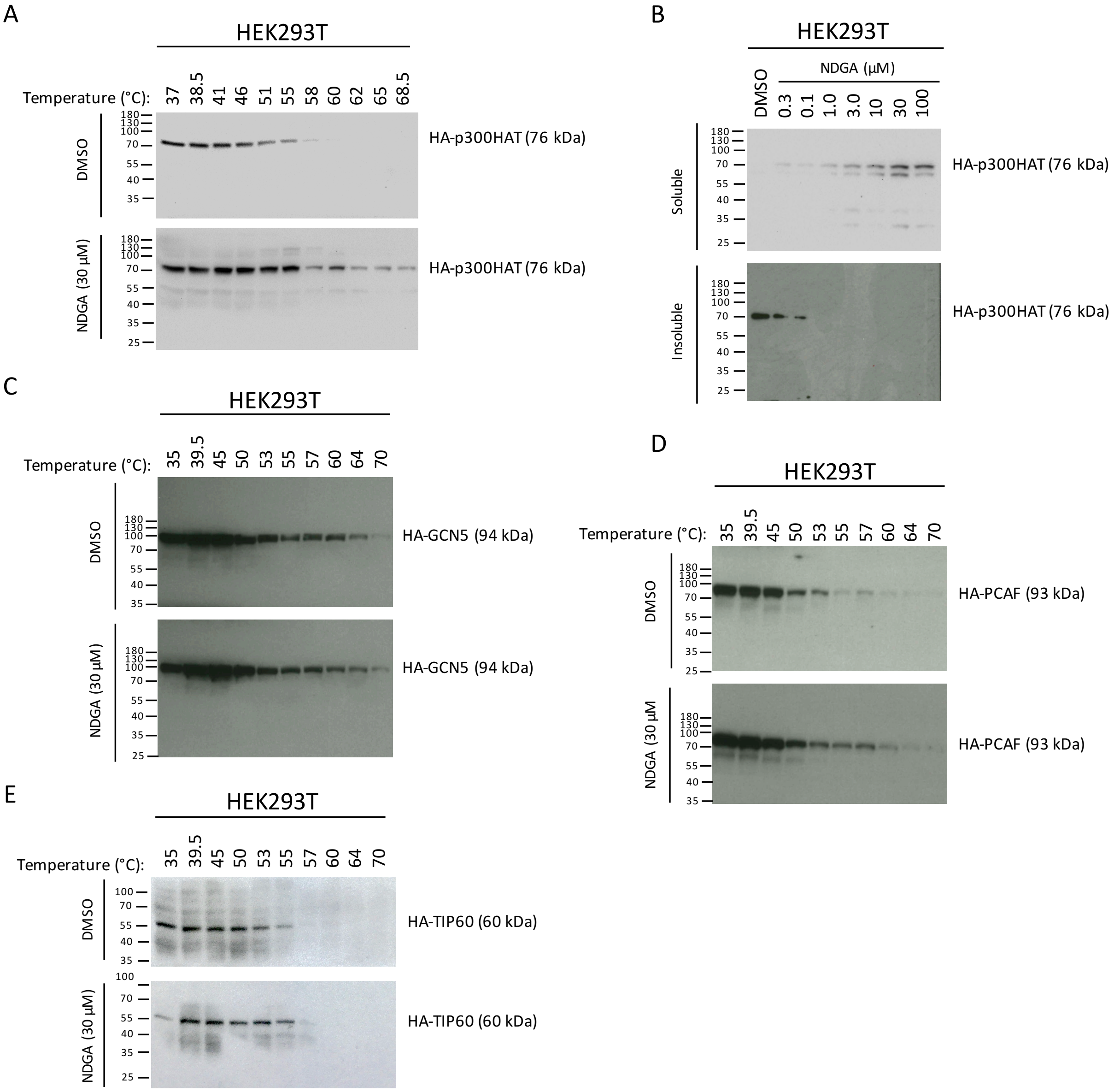

**Supplementary Figure 6**

A

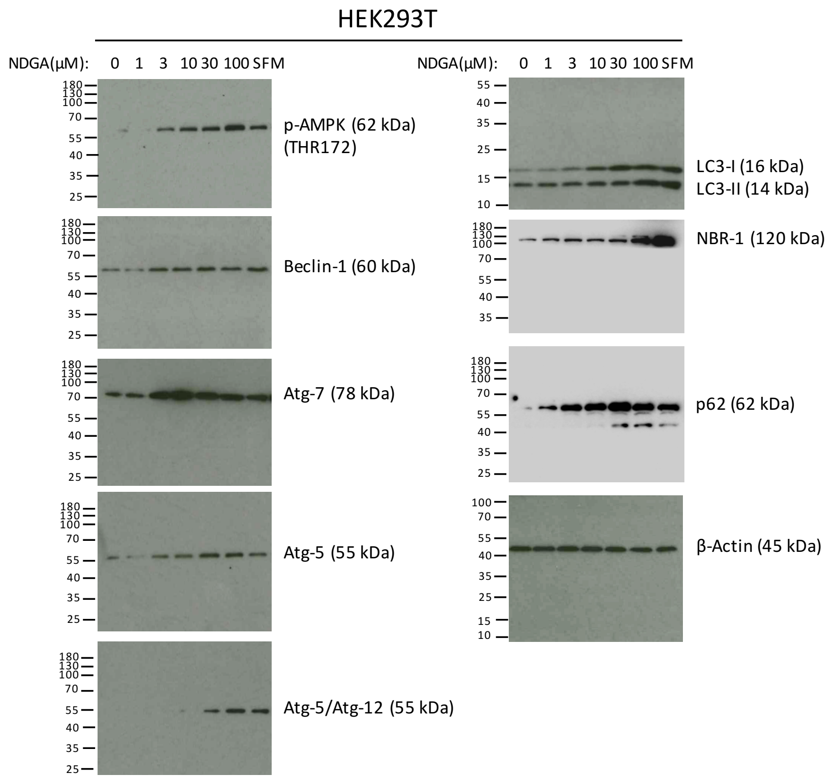


B

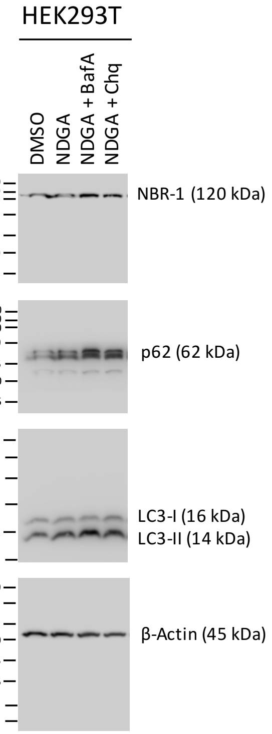


C

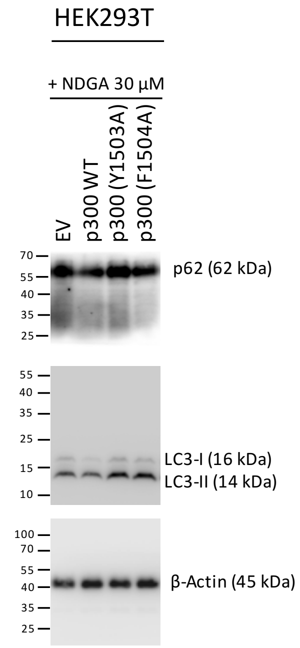


D

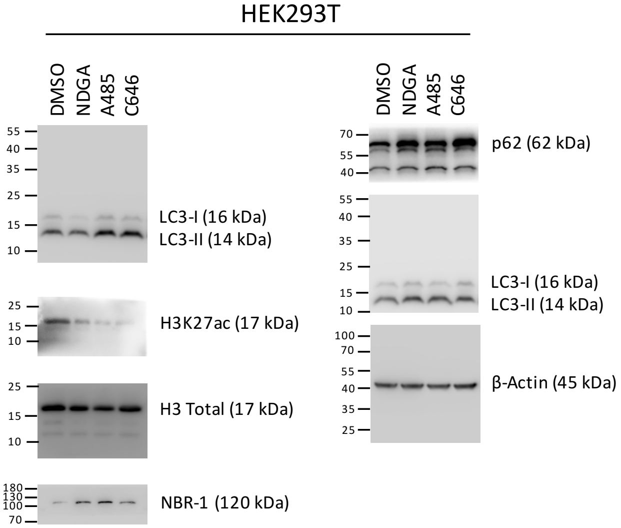

### Supplementary Figure 7

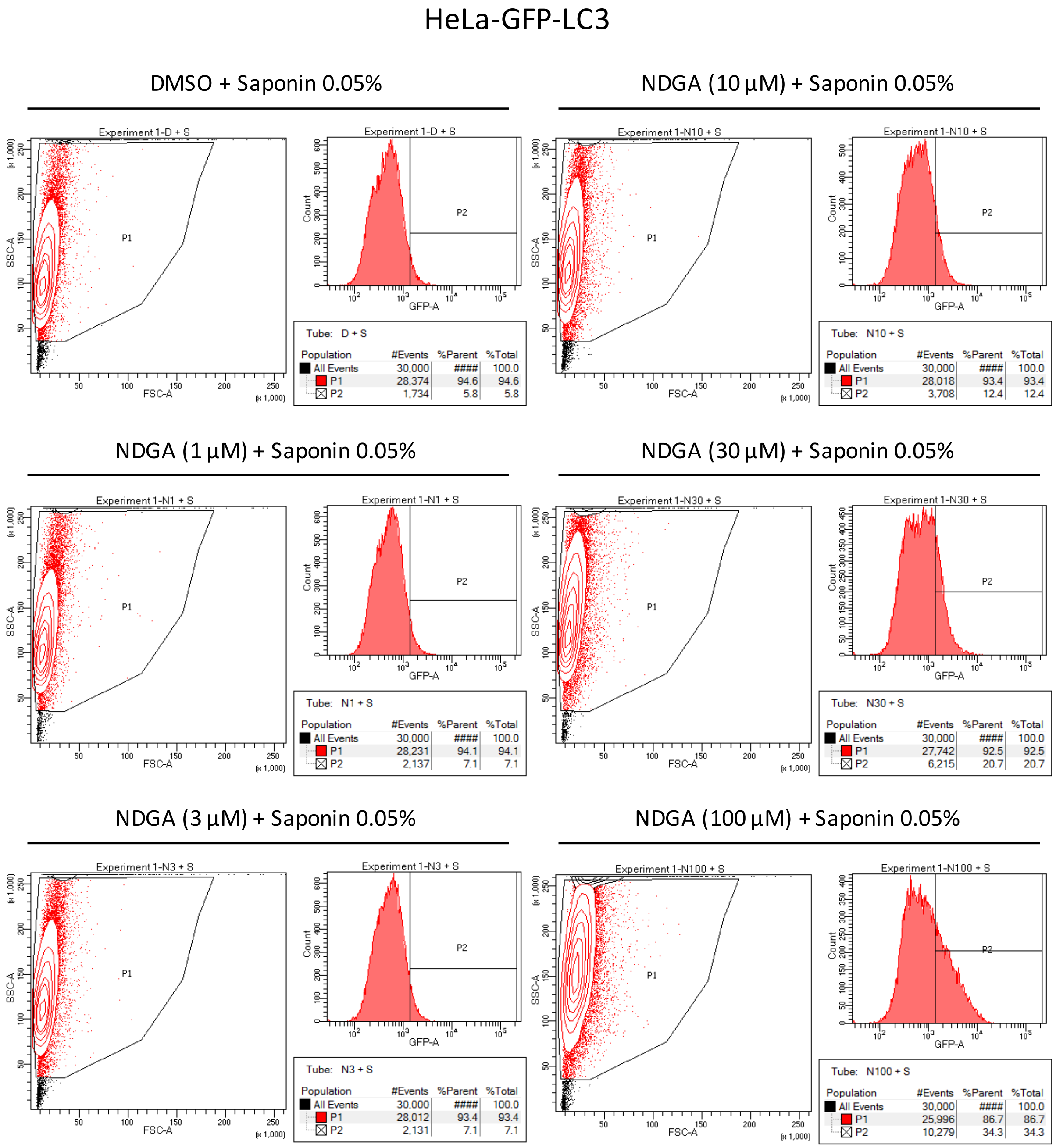


Tube: D + S

| Population | #Events | %Parent | %Total |
|---|---|---|---|
| All Events | 30,000 | #### | 100.0 |
| P1 | 28,374 | 94.6 | 94.6 |
| P2 | 1,734 | 5.8 | 5.8 |

Tube: N10 + S

| Population | #Events | %Parent | %Total |
|---|---|---|---|
| All Events | 30,000 | #### | 100.0 |
| P1 | 28,018 | 93.4 | 93.4 |
| P2 | 3,708 | 12.4 | 12.4 |

Tube: N1 + S

| Population | #Events | %Parent | %Total |
|---|---|---|---|
| All Events | 30,000 | #### | 100.0 |
| P1 | 28,231 | 94.1 | 94.1 |
| P2 | 2,137 | 7.1 | 7.1 |

Tube: N30 + S

| Population | #Events | %Parent | %Total |
|---|---|---|---|
| All Events | 30,000 | #### | 100.0 |
| P1 | 27,742 | 92.5 | 92.5 |
| P2 | 6,215 | 20.7 | 20.7 |

Tube: N3 + S

| Population | #Events | %Parent | %Total |
|---|---|---|---|
| All Events | 30,000 | #### | 100.0 |
| P1 | 28,012 | 93.4 | 93.4 |
| P2 | 2,131 | 7.1 | 7.1 |

Tube: N100 + S

| Population | #Events | %Parent | %Total |
|---|---|---|---|
| All Events | 30,000 | #### | 100.0 |
| P1 | 25,996 | 86.7 | 86.7 |
| P2 | 10,279 | 34.3 | 34.3 |

#### Supplementary Figure 8

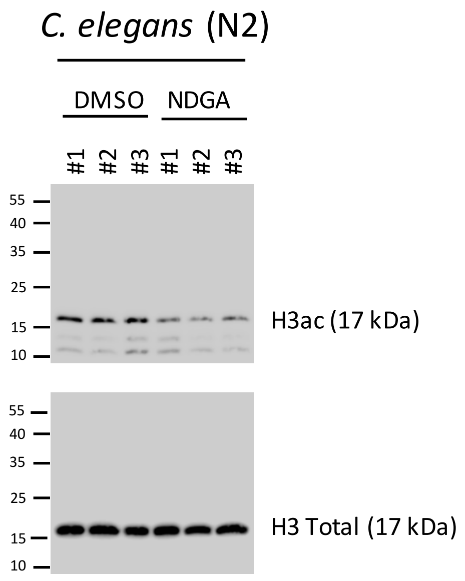

**Supplementary Table:** Correlation analysis for Fig. 2E

| **Pearson r** | **[NDGA] µM vs. 0 µg pCI-p300 (Full-length)** | **[NDGA] µM vs. 1 µg pCI-p300 (Full-length)** | **[NDGA] µM vs. 2 µg pCI-p300 (Full-length)** | **[NDGA] µM vs. 4 µg pCI-p300 (Full-length)** |
|---|---|---|---|---|
| r | -0.6809 | -0.8523 | -0.9269 | -0.9595 |
| R squared | 0.4636 | 0.7265 | 0.8591 | 0.9207 |
| P value | 0.1365 | 0.0311 | 0.0078 | 0.0024 |
| Significance (alpha = 0.05) | No | * | ** | ** |
| n | 6 | 6 | 6 | 6 |
| **Spearman r** | | | | |
| r | -1 | -0.9429 | -1 | -0.7714 |
| P value | 0.0028 | 0.0167 | 0.0028 | 0.1028 |
| Significance (alpha = 0.05) | ** | * | ** | No |
| n | 6 | 6 | 6 | 6 |

**Supplementary Notes**

Cloning (primers)

The HAT domain of human p300 protein was PCR amplified from pCi-p300 plasmid using p300HAT forward 5′-ATTTTCAAACCAGAAGAACTACGAC and p300HAT reverse 5′-GTCCTGGCTCTGCGTGTG; HA tag and stop codon were added by using HA-p300HAT forward 5′-ATGTACCCATACGATGTTCCAGATTACGCTATTTTCAAACCAGAAGAACTACG and p300HAT-stop reverse 5′-TAAGTCCTGGCTCTGCGTGTG primers. Final amplicon was cloned into pcDNA3.1(-) mammalian expression plasmid by Gibson Assembly kit (NEB) using HA-p300HAT-stop forward 5′-CGGCCGCCACTGTGCTGGATATGTACCCATACGATGTTCCAG and HA-p300HAT-stop reverse 5′-TGTGGTGGAATTCTGCAGATTAAGTCCTGGCTCTGCGTG primers. All constructs were validated by sequencing.

Antibodies

Primary antibodies anti-HA (Sigma, SAB1305536), anti-AMPKα (phospho-T172) (CST, 2535), anti-Beclin1 (CST, 3495), anti-LC3A/B (CST, 12741), anti-ATG5 (CST, 12994), anti-ATG7 (CST, 8558), anti-ATG12 (CST, 4180), anti-p62 (CST, 8025), anti-NBR1 (CST, 9891), and anti-β-actin (Sigma, A5441) were used for immunoblotting of total proteins. Primary antibodies anti-histone H2A (acetyl K5) (Abcam, ab1764), anti-histone H2A (acetyl K9) (Abcam, ab47816), anti-histone H2A (acetyl K13) (Abcam, ab177316), anti-histone H2A (Millipore, 07-146), anti-histone H2B (acetyl K5) (CST, 12799), anti-histone H2B (acetyl K11) (Abcam, ab40975), anti-histone H2B (acetyl K12) (CST, 5410S), anti-histone H2B (acetyl K15) (CST, 9083), anti-histone H2B (acetyl K16) (Abcam, ab40977), anti-histone H2B (acetyl K20) (CST, 2571S), anti-histone H2B (acetyl K24) (Abcam, ab176429), anti-histone H2B antibody (Abcam, ab1790), anti-histone H3 (acetyl K4) (Millipore, 07-539), anti-histone H3 (acetyl K9) (Millipore, 06-942), anti-histone H3 (acetyl K14) (CST, 7627), anti-histone H3 (acetyl-K18) (CST, 13998), anti-histone H3 (acetyl K27) (Millipore, 07-360), anti-histone H3 (acetyl K56) (CST, 4243), anti-histone H3 (tri-methyl K27) (Millipore, 07-449), anti-histone H3 antibody (Millipore, 07-690), anti-histone H4 (acetyl K5) (CST, 8647), anti-histone H4 (acetyl K8) (CST, 2594S), anti-histone H4 (acetyl K16) (Millipore, 06-762), anti-histone H4 antibody (Millipore, 07-108), anti-histone H3 (pan acetyl) (Millipore, 06-599) were used for immunoblotting of histones. Goat anti-rabbit-IgG-HRP (CST, 7074) and horse anti-mouse IgG-HRP (CST, 7076) were used as secondary antibodies in all immunoblotting experiments.

Flow Cytometry (gating)

Using the FSC/SSC, debris was removed by gating on the main cell population following saponin permeabilization. Positivity threshold for each group was defined on the basis of mock-treated (DMSO) sample. Identical positivity threshold was applied to all samples.
